# Decrease in Purifying Selection Pressures on Wheat Homoeologous Genes: Tetraploidization *vs* Hexaploidization

**DOI:** 10.1101/2024.04.07.587660

**Authors:** Akihiro Ezoe, Daisuke Todaka, Yoshinori Utsumi, Satoshi Takahashi, Kanako Kawaura, Motoaki Seki

**Author notes:** Corresponding authors Motoaki Seki; Akihiro Ezoe.

## Abstract

A series of polyploidizations in higher-order polyploids is the main event affecting the gene contents in a genome, and this is frequently observed in domesticated plants. Each polyploidization event is expected to lead to functional divergence because of the associated decrease in the selection pressures on the duplicated genes, but it is unclear whether the initial tetraploidization or the subsequent higher-order polyploidization has a greater evolutionary impact on the duplicated genes. To address this uncertainty, we focused on the *Triticum*–*Aegilops* complex lineage and compared the selection pressures before and after the tetraploidization and hexaploidization events. The results indicated that while both events decreased the selection pressures on homoeologous gene pairs (compared with the selection pressures on their ancestral diploid and tetraploid orthologous genes), the initial tetraploidization had a greater impact on the selection pressures on homoeologous gene pairs than the subsequent hexaploidization. This was supported by the analyzed expression patterns. Surprisingly, the decreases in the selection pressures on these homoeologous genes were independent of the existence of in-paralogs within the same subgenome. This result suggests that unique functions are maintained in the homoeologous genes, including the functions that are unlikely to be preserved in duplicate gene pairs derived from other duplication mechanisms. We also revealed their unique functions were different between the tetra- and hexaploidization (e.g., Reproductive system and chromosome segregation processes). The findings of this study imply that the substantial number of gene pairs resulting from multiple allopolyploidization events, especially the initial tetraploidization, may have been a unique source of functional divergence.

## Background

Duplicate genes are abundantly present in plant genomes. Immediately after gene duplication, duplicate genes possess redundant functions, the selection pressure on the gene pairs relaxes, leading to functional divergence between a pair of the duplicate genes [30, 50]. Divergent functions in duplicate genes are frequently observed and retained in most plant clades, and gene duplication is a prime source of genetic diversity in plants [37, 78]. Among the types of gene duplications, multiple rounds of polyploidization events in higher-order polyploids supply the most duplicate genes and are now recognized as a central diversifying force of plant gene functions [7, 31, 62]. Especially, higher-order polyploids are prevalent in domesticated plants, and the functional diversity of duplicate genes from a series of polyploidization events may also be important in the domesticated plants [19, 28, 40, 63].

In a series of polyploidizations, the tetraploidization of a diploid is the first step, and the relative scale of the impact on all genes, including singleton genes, is greater than that of higher-order polyploidizations. However, compared with tetraploidization events, hexaploidization or higher-order polyploidizations result in more duplicate genes with fewer evolutionary constraints. Therefore, the evolutionary effects of tetraploidization and hexaploidization or higher-order polyploidization events are expected to differ. Moreover, the consequences of each polyploidization event vary in a lineage-specific manner [32, 54, 73] [e.g., cotton and tobacco [15, 54]]. This highlights the unique consequences of each polyploidization event and suggests each step of multiple polyploidization events must be investigated independently. Unfortunately, this approach has not been employed in previous studies because of the large genomes of the analyzed species, a lack of comprehensive gene annotations, and the unknown order of polyploidization events [1, 23, 67]. Thus, the distinct effects of each step of multiple rounds of polyploidization events on functional divergence remain unclear, even though this type of polyploidization represents the main evolutionary mechanism underlying the massive induction of functional divergence and is an important source of genetic diversity in domesticated plants [53, 68]. The current study was conducted to compare tetraploidization events with hexaploidization or higher-order polyploidization events in terms of their evolutionary effects on functional divergence.

Hexaploid wheat is the most widely cultivated food crop because of recurring local adaptations on the basis of the genetic diversity due to two rounds of allopolyploidization, which is a type of polyploidization involving an inter-specific hybridization and chromosome doubling [23, 39]. The tetraploidization event, which occurred approximately 0.8 million years ago, involved the hybridization between the diploid progenitors of the A and B subgenomes (*Triticum urartu* and the closely related species *Aegilops speltoides*, respectively), whereas the hexaploidization event, which occurred 8,500–9,000 years ago, involved the hybridization between the tetraploid progenitor and the diploid progenitor of the D subgenome (*Triticum turgidum* and *Aegilops tauschii*, respectively) [38]. This *Triticum*–*Aegilops* complex lineage contains species with different ploidy levels, which is ideal for studying the effects of varying ploidy levels on the functional divergence of genes [75]. Moreover, there is no other allohexaploid organism whose genome information and gene annotations are available. Because of the rigorous mechanism mediating chromosome segregation in wheat, the karyotype and gene loci in each subgenome are highly conserved [27, 74]. The gene pairs located at the same locus, but originating from allopolyploidization events, are referred to as homoeologous genes [26]. This helps us distinguish the origins of each pair. Accordingly, wheat is the best species to compare the evolutionary impact on selection pressures between tetraploidization and hexaploidization events by focusing on their homoeologous gene pairs.

Earlier studies investigated the differences in protein functions and expression patterns following allopolyploidization events (e.g., subgenome dominance) [16, 17, 29, 54]. Some studies showed that the divergence in the expression patterns of homoeologous genes or the functions of the encoded proteins contributes to the acquisition of agronomic traits, including a high thousand-grain weight, long kernels, and anthocyanin pigmentation [16, 36, 45]. Similar to these earlier studies, a comparison of homoeologous gene pairs in terms of their expression patterns and the encoded protein functions can clarify the effects of allopolyploidization on tetraploids, but in hexaploids, the effects of each allopolyploidization event are indistinguishable. Hence, the effects of two distinct allopolyploidization events on selection pressures have not been directly compared. Furthermore, the presence of paralogs, which likely influence the functional divergence and decrease in the selection pressures on homoeologous pairs, has not been examined. In this study, we used recently updated genome information, including the assembly and annotation of the *Triticum*–*Aegilops* complex lineage with large genomes [4, 5, 41, 43, 76], to distinguish between allopolyploidization events and directly compare their effects on selection pressures. We also examined whether the presence of paralogs affects the decrease in selection pressures. Finally, we elucidated the functions of the homoeologous gene pairs under relaxed selection pressure after each allopolyploidization event. These analyses may reveal fundamental information regarding the functional diversification of homoeologous gene pairs following allopolyploidization events.

## Materials and Methods

### Annotation data of hexaploid wheat and progenitors

We obtained the nucleotide and encoded protein sequences of 41,505 genes in *T. urartu* (closest to the hexaploid wheat A subgenome) [41], 37,123 genes in *Ae. speltoides* (closest to the hexaploid wheat B subgenome) [4], 38,775 genes in *Ae. tauschii* (closest to the hexaploid wheat D subgenome) [43], 67,182 genes in *T. turgidum* ssp. *dicoccoides* (wild emmer wheat; tetraploid BBAA) [5], and 105,534 genes in *Triticum aestivum* cv. Chinese Spring DV418 (common wheat; hexaploid BBAADD) [76]. Only high-confidence sequences were used in our analyses.

### Definition of homoeologous gene pairs derived from the first and second allopolyploidization events (tetraploidization and hexaploidization) and inference of protein divergence rates

We obtained 45,702 sets of previously defined wheat homoeologous gene pairs [51]. Because the annotation versions were RefSeq-v1.0 or -v1.1 gene models, we updated them to RefSeq-v2.1 gene models [76]. Of these sets, 26,183 consisting of high-confidence genes were used. For each gene in a homoeologous gene set, we performed a reciprocal best-hit search (E-value < 1 × 10^−4^) of the genes in the ancestral A, B, and D genomes and in the allotetraploid A and B subgenomes using diamond-v2.1.6.160 [11], which resulted in the identification of 13,335, 17,157, 20,824, 19,451, and 19,314 orthologous genes, respectively, (Supplementary Table S1). In the two relationships in a homoeologous gene set related to the second allopolyploidization event (D-A and D-B), the A or B gene closer to the D gene was selected for each homoeologous gene. On the basis of this procedure, 16,258 homoeologous gene pairs between the tetraploid A and B subgenomes and 18,807 homoeologous gene pairs between the hexaploid A or B and D genes were defined as the homoeologous pairs derived from the first and second allopolyploidization events, respectively (Supplementary Table S1). To compare the evolutionary impact between the orthologous and homoeologous gene pairs, we focused on 9,164 (first round) and 13,997 (second round) gene sets consisting of conserved orthologous and homoeologous gene pairs.

Selection pressures on homoeologous and ancestral gene pairs were inferred by the pairwise alignment and file format conversion performed using mafft-v7.407 and pal2nal-v14 [35, 59]. The synonymous substitution rate (Ks) and non-synonymous substitution rate (Ka) for a gene pair were estimated using the yn00 module in PAML-v4.9 [71, 72]. To eliminate paralogous pairs from homoeologous and orthologous pairs, we focused on the gene sets in which Ks was less than 0.2 (according to the relationship between the time from duplication events and Ks) [14, 46, 48, 65]. We finally used 6,796 and 10,939 (A-D: 5,166, B-D: 5,773) gene sets for the first and second polyploidization events, respectively (Supplementary Table S1).

### Expression analysis

We obtained an expression atlas of hexaploid wheat in 850 conditions from an earlier study [51]. The annotation of this expression data was completed on the basis of the RefSeq-v1.0 and -v1.1 gene models. Hence, we updated the annotation to RefSeq-v2.1 gene models [76]. Among the 850 conditions, 161 were for gene models that cannot be converted to RefSeq-v2.1 gene models. Accordingly, we focused on the remaining 689 conditions. The expression conditions in this data were described previously [51]. For the 6,796 A-B and 10,939 A-D or B-D homoeologous pairs, we calculated the correlation coefficients for all expression patterns. These correlation coefficients were converted to reflect the divergence of the expression patterns according to the earlier study [21]. Additionally, by dividing by Ks, the expression pattern divergence rates were determined [21, 37] and used to represent the selection pressures on gene expression.

### Definition of in-paralogs

In-paralogs were defined as duplicated genes that exist within the same subgenome of hexaploid wheat. Because high similarity between homoeologous genes and in-paralogs was expected to influence selective pressure, in-paralogs were searched under the conditions: E-value < 1 × 10^−9^ and coverage > 75%. By this homology search, 14,285, 14,949, and 13,208 in-paralologs were identified in the A, B, and D subgenomes, respectively. For a single homeologous gene, we used labels as the presence or absence of in-paralogs, and this information is listed in Supplementary Table S1.

### Gene Ontology (GO) enrichment analysis

To characterize functional differences between tetraploidization and hexaploidization, the enriched GO terms were inferred using a Chi-squared test, and the false discovery rate (FDR) was used for correction. To focus on the functional differences between the two events, the background for this test was not the entire genome but all homoeologous genes. This analyses were conducted using Panther [47]. A total of 3,442 GO terms were related to at least one homoeologous gene, and among these, 224 significantly enriched GO terms (FDR < 0.05) were identified. All of these were listed in Supplementary Table S2.

## Results

### Evolutionary effects of two rounds of allopolyploidization events on the selection pressures for protein sequences

To examine the impact of each allopolyploidization event on selection pressures, we used Ka/Ks as an indicator of the selection pressure on orthologous and homoeologous gene pairs. We focused on the following five species: *T. urartu* (AA), *Ae. speltoides* (BB), *Ae. tauschii* (DD), *T. turgidum* (BBAA), and *T. aestivum* (BBAADD) (Fig. 1a). By comparing Ka/Ks between ancestral A-B (orthologous pairs before the first allopolyploidization) and *T. turgidum* A-B (homoeologous pairs after the first allopolyploidization) and between *T. turgidum* AB-ancestral D (orthologous pairs before the second allopolyploidization) and *T. aestivum* AB-D (homoeologous pairs after the second allopolyploidization) (Fig. 1a), we analyzed the decreases in selection pressures after the first and second allopolyploidization events in a pairwise manner between orthologous and homoeologous pairs. The Ka/Ks values were higher for the homoeologous gene pairs than for the orthologous gene pairs for the first and second allopolyploidization events (Fig. 1b and Supplementary Table S1; first: P = 3.40 × 10^−108^, second: P = 3.01 × 10^−6^, two-tailed paired Wilcoxon signed rank test). The homoeologous pairs under positive selection were also identified (Ka/Ks > 1 and orthologous < homoeologous; first and second allopolyploidization events: 46 and 66, respectively). These results indicated that the selection pressures relaxed after the first and second allopolyploidization events.

**Fig. 1:**
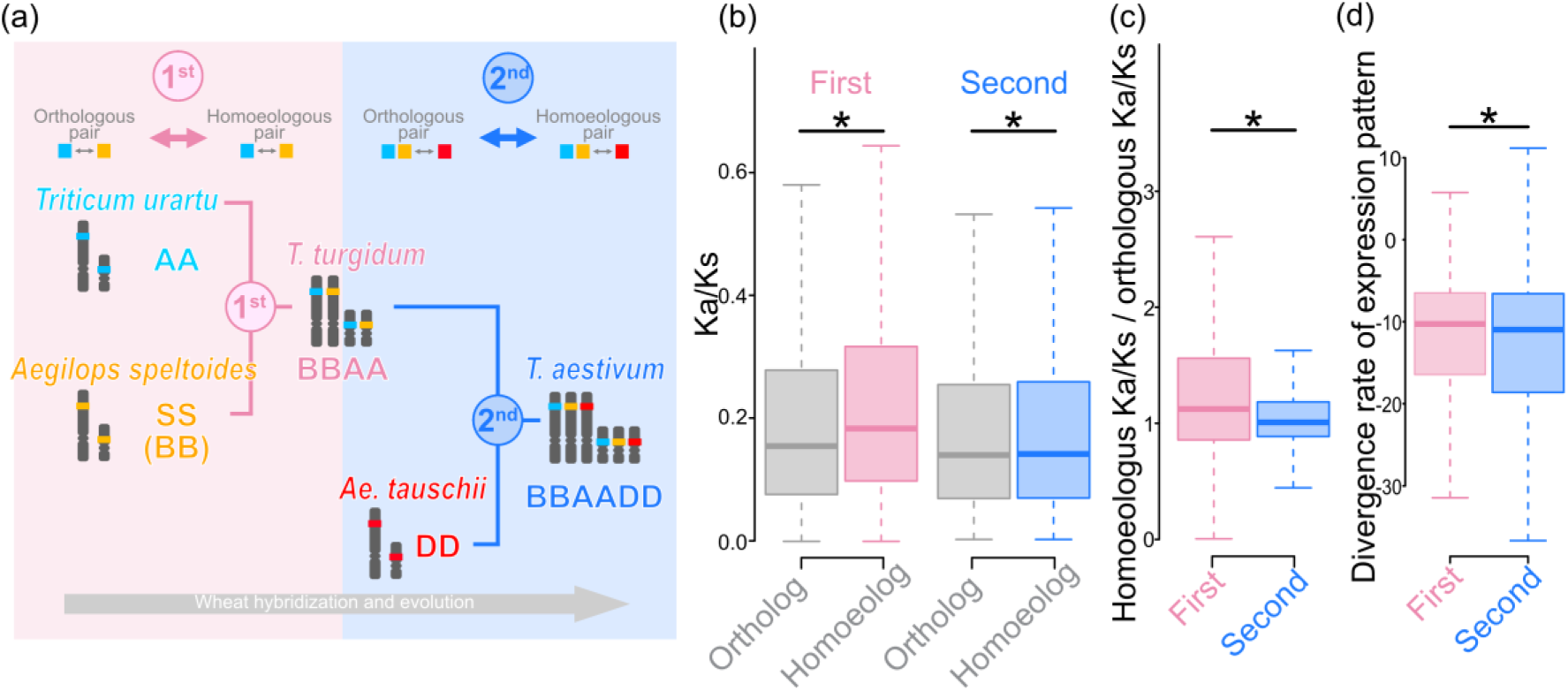
Two rounds of allopolyploidization events in the *Triticum*–*Aegilops* complex and differences in the selection pressures on protein sequences and expression patterns. (a) Flow of the first (tetraploidization) and second (hexaploidization) allopolyploidization events. The first event occurred between *Triticum urartu* and *Aegilops speltoides*, whereas the second event occurred between *Triticum turgidum* and *Aegilops tauschii*. Gray bidirectional arrows between an orthologous gene pair and between a homoeologous gene pair indicate the pairs used to infer selection pressures. Pink and blue bidirectional arrows indicate the pairs used to assess evolutionary effects on selection pressures after the first and second allopolyploidization events, respectively. (b) Distributions of Ka/Ks between orthologous and homoeologous gene pairs are presented in box plots with the solid horizontal line indicating the median value, the box representing the interquartile range (25%–75%), and the dotted line indicating the first to 99th percentile. The asterisk indicates a significant difference (P < 0.05, two-tailed paired Wilcoxon signed rank test). (c and d) Distributions of the ratios of Ka/Ks and the expression pattern divergence rates for the first and second allopolyploidization events are presented in box plots with the solid horizontal line indicating the median value, the box representing the interquartile range (25%–75%), and the dotted line indicating the first to 99th percentile. The asterisk indicates a significant difference (P < 0.05, two-tailed Wilcoxon rank sum test).

We subsequently directly compared the decreases in selection pressures between the first and second allopolyploidization events by focusing on the Ka/Ks ratios between the orthologous and homoeologous gene pairs. The ratios were significantly higher for the first allopolyploidization event (average: 1.68) than for the second allopolyploidization event (average: 1.25) (Fig. 1c and Supplementary Table S1; P = 4.93 × 10^−82^, two-tailed Wilcoxon rank sum test). This result indicated that the first allopolyploidization event had a greater impact than the second allopolyploidization event on the selection pressures on homoeologous gene pairs.

### Evolutionary effects of two rounds of allopolyploidization events on the selection pressures for expression patterns

The two rounds of allopolyploidization events had similar evolutionary effects on the protein sequences encoded by the homoeologous gene pairs and the corresponding expression patterns. The comparison of the expression pattern divergence rates of 6,796 and 10,939 homoeologous gene pairs for the first and second allopolyploidization events indicated that the expression pattern divergence rates were higher for the first event than for the second event (Fig. 1d and Supplementary Table S1; P = 1.04 × 10^−9^, two-tailed Wilcoxon rank sum test). These results implied that the selection pressures on the expression patterns were relaxed after the first event, which was consistent with the decreases in the selection pressures on the protein sequences.

### Homoeologous gene pairs under positive selection

Out of the genes under the relaxed selection pressures, some genes were expected to be under positive selection [12, 18]. Furthermore, it is known that young duplicate genes, like the wheat homoeologous pairs, are more frequently under positive selection [12]. Genes under positive selection underwent rapid functional changes contributing to the adaptation of the species, and tend to be specific functional groups of genes such as reproductive genes and immunity [18, 57]. Therefore, it was necessary to distinguish homoeologous gene pairs under positive selection from the other homoeologous gene pairs. We then categorized homoeologous pairs into three categories: those under relaxed selection pressure after allopolyploidization events (Ka/Ks: orthologous < homoeologous), those under positive selection (Ka/Ks: orthologous < homoeologous and Ka/Ks > 1), and others (Fig. 2a). For the first allopolyploidization event, we found 4,113, 46 and 2,637 pairs in these categories, respectively. For the second allopolyploidization event, we identified 5,064, 66 and 5,809 pairs in these categories, respectively. This categorization was used in the following analyses.

**Fig. 2:**
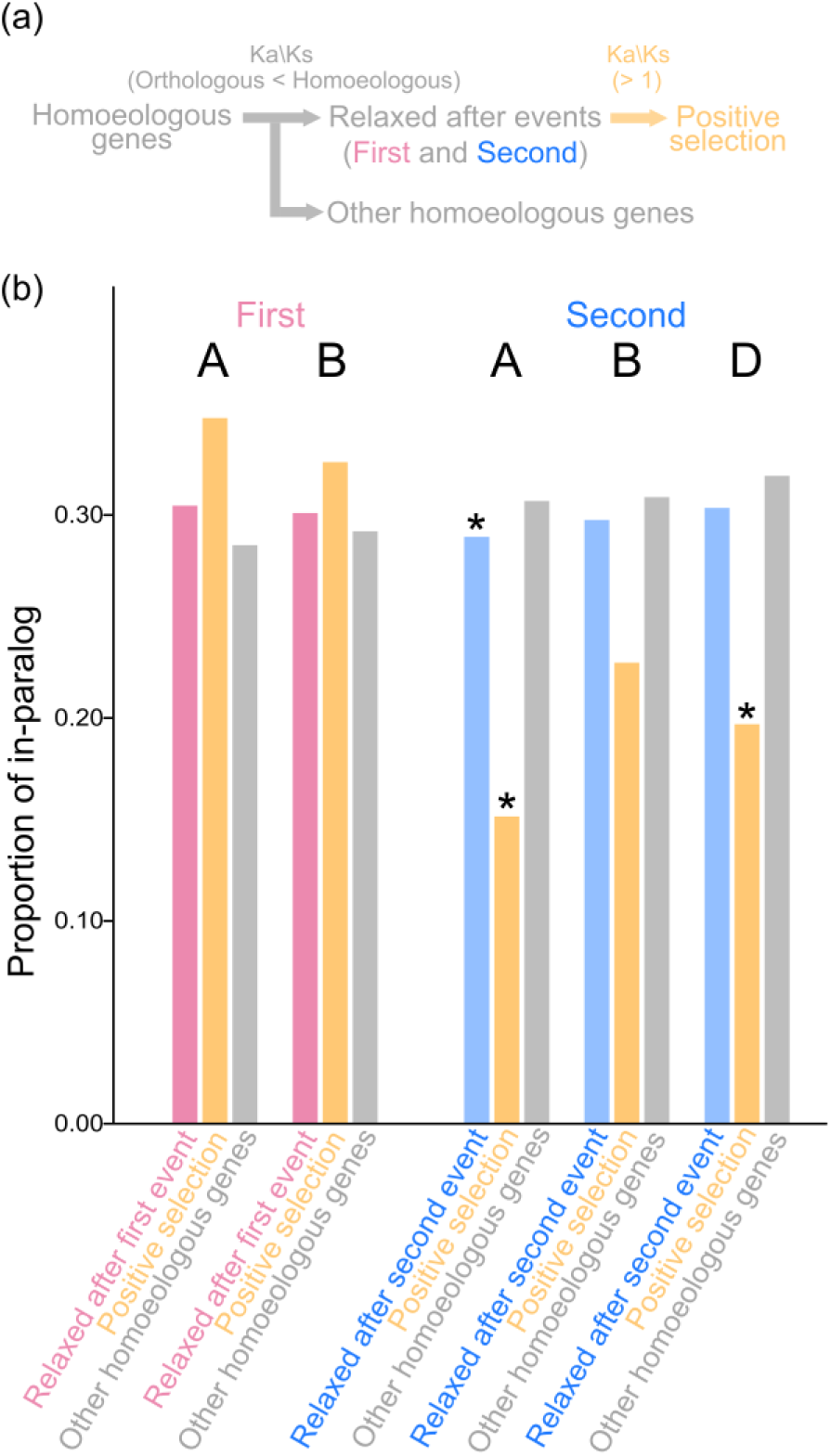
In-paralogs in homoeologous genes under relaxed selection after first and second events and positive selection. (a) Classification of homoeologous genes into those under relaxed selection after first and second events and positive selection. If a pair of homoeologous genes has higher Ka/Ks than their orthologous pairs, the pair is classified as a pair under relaxed selection pressure. Out of the pairs under relaxed selection pressure, if Ka/Ks is over 1, the pair is classified as a pair under positive selection. (b) Proportions of the homoeologous gene pairs with and without in-paralogs for each subgenome. The asterisk indicates a significant difference (P < 0.05, Chi-squared test).

### The decrease in the selection pressures on homoeologous gene pairs occurred independently of the existence of paralogs within each subgenome

The decrease in the selection pressure on homoeologous gene pairs might be influenced by the existence of paralogous genes. To assess this possibility, we compared the proportions of the homoeologous gene pairs with in-paralogs (paralogous genes in the same subgenome) in the above three categories (Fig. 2a). We examined the proportions of the homoeologous gene pairs with and without in-paralogs for each subgenome. As a result, there was no case where the proportions of homoeologous genes with in-paralogs under relaxed selection pressure was significantly higher than those without in-paralogs (Fig. 2b and Supplementary Table S3). Therefore, the decrease in selection pressures may have been unaffected by the duplicated genes derived from other mechanisms (e.g., tandem duplication). The results also indicated that for the second allopolyploidization event, homoeologous genes under relaxed selection pressure and those under positive selection tended to lack in-paralogs in the A and D subgenomes (Fig. 2b and Supplementary Table S3; P = 9.13 × 10^−3^ and P = 4.43 × 10^−2^, Chi-squared test), respectively. Furthermore, in the A subgenome, the second allopolyploidization event-derived homoeologous genes under relaxed selection pressure tended to lack in-paralogs (Fig. 2b and Supplementary Table S3; P = 4.45 × 10^−2^, Chi-squared test). Accordingly, these homoeologous pairs might be a unique group of genes exclusively duplicated by allopolyploidization.

### Enrichment of Gene Ontology (GO) terms among the homoeologous genes under relaxed selection pressure or under positive selection

To investigate the unique functions among the homoeologous genes, a GO enrichment analysis was conducted for the homoeologous genes under relaxed selection pressure or positive selection and the other homoeologous genes derived from the first and second allopolyploidization events.

For the first allopolyploidization event-derived homoeologous genes under relaxed selection pressure, there were 43 over-represented GO terms (Supplementary Table S2). These GO terms were related to reproductive system development [e.g., reproductive structure development (GO:0048608) and developmental process involved in reproduction (GO:0003006)] and abiotic stress responses [e.g., cellular response to abiotic stimulus (GO:0071214) and cellular response to environmental stimulus (GO:0104004)] (Fig. 2). The homoeologous genes annotated with the GO terms related to reproductive system development included 12 genes encoding MADS-box transcription factors, which is consistent with earlier research that detected changes in gene expression patterns immediately after an allopolyploidization event as well as large-scale duplication events involving these transcription factor genes [2, 69]. Therefore, it is reasonable that the selection pressures on the genes related to the reproductive system relaxed after the first allopolyploidization event. Furthermore, a previous study showed that genes related to abiotic stress responses tended to be retained after allopolyploidization events [16]. The functional diversity among the homoeologous genes related to abiotic stress responses may have contributed to the considerable expansion of the wheat cultivation area.

For the first allopolyploidization event-derived homoeologous genes under positive selection, there were two over-represented GO terms (Supplementary Table S2). The GO term with the lowest FDR was unclassified (GO:UNCLASSIFIED) (Fig. 2). Considering the first allopolyploidization event occurred in wild environments [34], it is possible that the unclassified processes represent previously undiscovered factors contributing to the establishment of allopolyploids in wild environments. These unclassified genes should be the focus of future investigations.

There were 114 over-represented GO terms assigned to the second allopolyploidization event-derived homoeologous genes under relaxed selection pressure (Supplementary Table S2). These GO terms were related to the regulation of pH [e.g., monoatomic ion transport (GO:0006811) and pH regulation (GO:0006885)], DNA repair [e.g., interstrand cross-link repair (GO:0036297) and double-strand break repair via homologous recombination (GO:0000724)], and chromosome segregation [e.g., chromosome segregation (GO:0007059) and spindle assembly (GO:0051225)] (Fig. 2). The homoeologous genes associated with the regulation of pH included Na or K transporter genes such as high-affinity K^+^ transporter (HKT) family members. The homoeologous genes under relaxed selection pressure included 11 HKT genes. An earlier study showed that salt tolerance differed between tetraploid progenitors and newly synthesized hexaploids, but also between newly synthesized hexaploids and natural hexaploids; the salt tolerance was due to HKTs [70]. This indicates that wheat HKT gene functions change after the second allopolyploidization event, consistent with our findings. Moreover, the regulation of DNA repair and chromosome segregation processes is altered by allopolyploidization events [9], which is also consistent with our findings.

The second allopolyploidization event-derived homoeologous genes under positive selection were annotated with 65 over-represented GO terms (Supplementary Table S2), including those related to gene silencing [e.g., DNA methylation-dependent heterochromatin formation (GO:0006346) and gene silencing by RNA-directed DNA methylation (GO:0080188)], guard cell differentiation [e.g., stomatal complex morphogenesis (GO:0010103) and guard cell differentiation (GO:0010052)], meiosis [e.g., meiosis II cell cycle process (GO:0061983) and gamete generation (GO:0007276)], and DNA repair [e.g., interstrand cross-link repair (GO:0036297) and double-strand break repair via nonhomologous end joining (GO:0006303)] (Fig. 2). Notably, gene silencing by methylation and meiosis is essential for stabilizing polyploid plants via the elimination of evolutionary constraints regarding excessive gene dosage and expression as well as chromosome segregation [25, 49, 52], which is consistent with our findings. The over-representation of GO terms associated with guard cell differentiation suggests that these genes may contribute to the physiological adaptations of polyploid plants, potentially by improving their water use efficiency and gas exchange capabilities. In fact, a recent study showed that guard cell size and gas exchange capabilities tend to increase immediately after allopolyploidization events [64]. Therefore, it is reasonable that the functions of the genes related to guard cell differentiation diversified after the allopolyploidization event.

Overall, the enriched GO terms among the homoeologous gene pairs under relaxed selection pressure or positive selection were related to genome stability, including DNA repair, chromosomal segregation, gene silencing, and meiosis. This indicates that regulatory changes that maintain genome stability via homoeologous genes are altered after every allopolyploidization. This is in accordance with previous research that demonstrated the importance of genome stability for establishing allopolyploids [9, 22]. Additionally, given that these genes tend to be lost after whole-genome duplication events [3, 8], the functional divergence between homoeologous gene pairs due to positive selection is reasonable. Moreover, the retention of these homoeologous genes may be a unique feature. Fig. 3

**Fig. 3:**
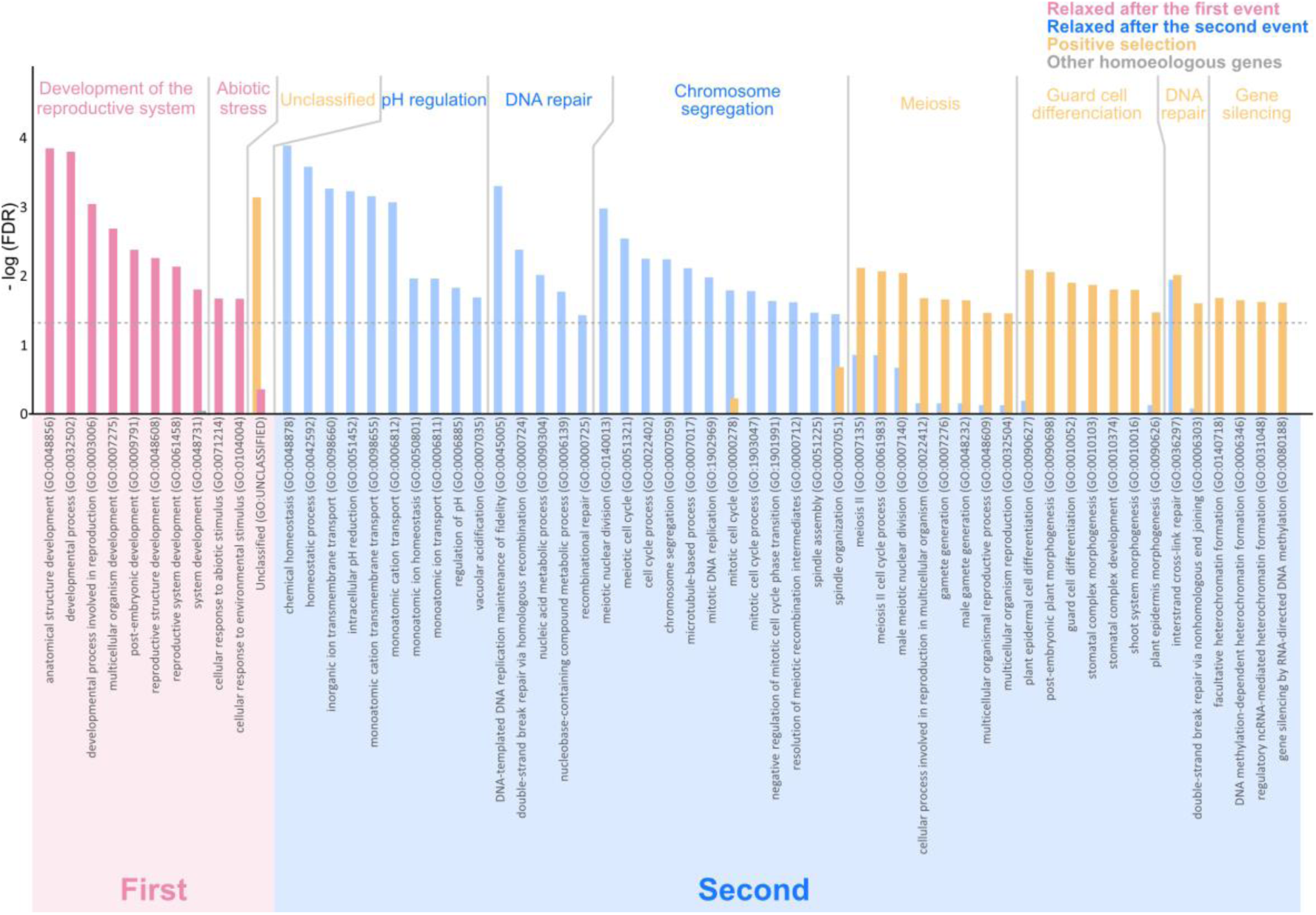
Enriched functional categories among the homoeologous genes. The degree of enrichment of each GO term is represented using the minus logarithm of the false discovery rate (FDR) inferred by a Chi-squared test. The dotted line indicates the criterion of FDR < 0.05. The x-axis presents the enriched GO terms, with gray lines separating broad classifications of functional categories. As indicated in the upper right part of the figure, the pink/blue, yellow, and gray bars represent the homoeologous genes under relaxed selection pressure or positive selection and the other homoeologous genes.

## Discussion

Higher-order polyploids have undergone multiple rounds of polyploidization events, resulting in significant genetic diversity [7, 31, 62]. According to previous studies, the genetic diversity of hexaploid wheat, which was due to tetraploidization and hexaploidization events, contributed to the cultivation of wheat [6, 19, 24]. However, tetraploidization and hexaploidization events are believed to have distinct evolutionary effects on the functional diversity of wheat genes because tetraploidizations have a greater effect on the entire genome, including singleton genes, than hexaploidizations, but hexaploidizations result in a greater abundance of duplicated genes with fewer evolutionary constraints. In this study, we compared the effects of tetraploidization and hexaploidization events to determine which event is more responsible for the decreases in the selection pressures that led to genetic diversity and the resulting functional divergence of homoeologous genes. To compare the effects of allopolyploidization events on selection pressures, we analyzed the fold-changes to the selection pressures between the ancestral and tetraploid or hexaploid wheat plants. First, we confirmed that selection pressures on the homoeologous gene pairs were relaxed after the tetraploidization and hexaploidization events. Moreover, the homoeologous gene pairs under positive selection were also identified. Second, we observed that the tetraploidization event influenced the selection pressures more than the hexaploidization event. This observation was supported by the results reflecting the relaxed selection pressure on expression patterns. These findings indicate that homoeologous gene pairs contributed to the functional divergence after both allopolyploidization events and that the tetraploidization event tended to have a more substantial impact on protein sequences and expression patterns than the hexaploidization event. The tetraploidization and hexaploidization events may have distinct evolutionary effects on selection pressures and the functional divergence of homoeologous genes.

The existence of paralogous genes is one of the factors associated with relaxed selection pressures [60]. This compelled us to investigate the evolutionary relationship between allopolyploidization and other duplication mechanisms. The homoeologous pairs under relaxed selection pressures were not affected by the existence of in-paralogs, but rather the homoeologous gene pairs under relaxed selection pressure or positive selection in subgenomes A and D lacked in-paralogs. This indicates that the homoeologous pairs tended to have functions that differed from the functions of the duplicate genes derived from other duplication mechanisms. To explore their functions, we performed a GO term enrichment analysis. The genes under relaxed selection pressure derived from the first allopolyploidization event were associated with the development of the reproductive system and abiotic stress responses, suggestive of their contributions to the domestication of wheat. In contrast, the enriched GO terms for the second allopolyploidization event were related to pH regulation, chromosome segregation, and DNA repair. Additionally, the functions of the genes under positive selection were related to meiosis, DNA repair, methylation, and guard cell differentiation, implying the homoeologous genes helped maintain polyploid stability and enhanced the adaptations to physiological and genomic changes caused by allopolyploidization. The enriched GO terms assigned to the homoeologous genes under relaxed selection pressure and positive selection were related to chromosome segregation, meiosis, and DNA repair. Given that these genes tended to be lost immediately after other duplication events, relaxed or positive selection may be important for the functional divergence of homoeologous gene pairs necessary to prevent the deleterious effects of gene dosage, which is unique to homoeologous genes [8, 50, 55]. Another study showed that the loss of homoeologous genes related to chromosome segregation and meiosis was lethal to tetraploid yeast [58]. Moreover, wheat homoeologous genes with similar functions are reportedly conserved [3]. These results supported our findings. Hence, we speculate that the functional divergence of the genes involved in chromosome segregation, DNA repair and meiosis processes might be an essential part of allopolyploidization events. In addition, considerable changes to the functions of these genes may have helped prevent the harmful effects of allopolyploidization events. This is a potentially important insight into the functional divergence following allopolyploidization. The GO enrichment analysis provided key insights into the functional diversification of homoeologous genes and evolutionary adaptations after allopolyploidization events.

Allopolyploids have two types of functional diversities through species-specific genes acquired after the divergence of the common ancestors and via the relaxed selection pressures on homoeologous gene pairs [16]. Although previous studies examined the functional diversity in ancestral species-specific genes (e.g., the diversity due to habitat expansion after tetraploidization events) between tetraploids and their diploid ancestors [23, 56], the functional diversity following the relaxation of selection pressures by allopolyploidization events remains unexplored. Most of the relevant research focused on the gain and loss of genes in ancestral diploid species; the results of these studies reflect the lack of dominance among the A, B, and D subgenomes [33, 54, 66]. The present study showed that the tetraploidization and hexaploidization events led to relaxed selection pressures. Furthermore, their evolutionary effects on selection pressures differed, suggestive of the importance of focusing on ancestral species-specific functions as well as homoeologous gene pairs derived from each allopolyploidization event.

Although generating and analyzing knock-out mutants is one of the best approaches to clarifying gene functions, its applicability for hexaploid wheat is somewhat limited by the considerable redundancy of homoeologous genes [20, 38], which has been a major obstacle in wheat breeding programs [10]. The current study potentially identified functionally divergent homoeologous gene pairs that were likely associated with morphologically abnormal phenotypes because the homoeologous gene pairs under relaxed or positive selection may not compensate for the loss of a functional counterpart [21, 42]. Thus, the analysis of knock-out mutants in the homoeologous gene pairs under relaxed or positive selection may be effective. Both TraesCS7A03G0884000 and TraesCS7B03G0730800 were identified as candidates for mutant analyses. These genes were highly conserved in the diploid ancestors, but TraesCS7A03G0884000 was under positive selection after the tetraploidization. This gene was annotated as an oligosaccharyltransferase (OST) subunit, which is crucial for N-linked glycosylations [70]. The OST complex is composed of several subunits, each contributing to the overall enzyme function and specificity. A recent study showed that STAUROSPORINE AND TEMPERATURE SENSITIVE3 (STT3) in hexaploid wheat is an OST subunit that helps regulate plant innate immunity and grain weight [77]. Therefore, the positive selection of TraesCS7A03G0884000 may enhance immunity and increase the grain weight of wheat. Among the other candidates, TraesCS4D03G0720800 and TraesCS4A03G1013900 were identified as a homoeologous pair under positive selection after hexaploidization. These genes were highly conserved in the diploid and tetraploid ancestors, but TraesCS4D03G0720800 was under positive selection after the hexaploidization event. This gene encodes flowering-promoting factor 1-like protein 5, which is a critical factor that regulates flowering time as a common downstream target of SOC1, VRN1, and phytochromes [44]. In fact, our data suggest that the homoeologous gene pair for phytochrome B (TraesCS4D03G0456400), which is one of the master regulators of photoperiodic processes [61], also experienced a decrease in the purifying selection after the hexaploidization event. Conversely, there was no decrease in the purifying selection of the orthologous gene pair nor after the tetraploidization event. Thus, the photoperiodic regulation of the flowering time may have been modified after the hexaploidization event as preceding studies showed [13]. However, these genes have not been sufficiently functionally annotated and a forward genetics research approach has not been applied. These genes may be functionally characterized relatively simply by mutating either the A or B subgenome of hexaploid wheat and then examining the mutants for phenotypic changes.

## Conclusions

In this study, we revealed that tetraploidization events decreased the selection pressure more than hexaploidization events. In addition, the homoeologous genes were typically unaffected by the existence of paralogs, suggesting functions that may not be preserved in duplicate gene pairs produced by other duplication mechanisms are maintained by homoeologous gene pairs. We also elucidated the unique functional tendencies of the homoeologous gene pairs such as genomic stability and abiotic stress response. Therefore, the substantial abundance of homoeologous gene pairs generated by multiple allopolyploidization events may have been critical for the functional divergence contributing to the establishment of allopolyploids and the adaptations to new environmental conditions during the cultivation of wheat. Future research will need to focus on species-specific functions of ancestral species as well as functionally diverse homoeologous gene pairs derived from individual allopolyploidization events. The findings of such research may provide valuable insights relevant to studies on the complex allo-/auto- and higher-order polyploidizations that occurred during the evolution of plants.

## Contributions

A.E. and M.S. conceived the study. M.S. and K.K. supervised the project. A.E., D.T., Y.U., and T.S. processed the sequencing and expression data. A.E. performed the evolutionary analyses. All authors discussed the results and wrote the manuscript. All authors read and approved the final manuscript.

## Acknowledgments

We thank Kentaro Yoshida, Kazumasa Shirai, and Kousuke Hanada for providing useful suggestions regarding this study. This research was supported by a grant from the RIKEN Center for Sustainable Resource Science (CSRS) (to M.S.), the Incentive Research Grant in RIKEN, Grant-in-Aid for Early-Career Scientists (23K13932) (to A.E.), the RIKEN Special Postdoctoral Researcher Program (to A.E.), the Teijin-Kumura scholarship (to A.E.), and RIKEN research funding for researchers returning to work/research after an interruption caused by life events (to A.E.). We thank Edanz (https://jp.edanz.com/ac) for editing a draft of this manuscript.

## Conflict of Interest Statement

All authors have no conflicts of interest to declare.

## Supplementary Information

Supplementary Table S1. Information regarding homoeologous genes, orthologous genes, paralogs, and selection pressures

Supplementary Table S2. Enriched Gene Ontology terms, over/under-representation, and false discovery rates

Supplementary Table S3. Number of homoeologous gene pairs with and without in-paralogs

## Notes

### Competing Interest Statement

The authors have declared no competing interest.

